# Associations of coffee genetic risk scores with coffee, tea and other beverages in the UK Biobank

**DOI:** 10.1101/096214

**Authors:** Amy E. Taylor, Marcus R. Munafò

## Abstract

**Background:** Genetic variants which determine amount of coffee consumed have been identified in genome-wide association studies (GWAS) of coffee consumption; these may help to further understanding of the effects of coffee on health outcomes. However, there is limited information about how these variants relate to caffeinated beverage consumption more generally.

**Aims:** To improve phenotype definition for coffee consumption related genetic risk scores by testing their association with coffee, tea and other beverages.

**Methods:** We tested the associations of genetic risk scores for coffee consumption with beverage consumption in 114,316 individuals of European ancestry from the UK Biobank. Drinks were self-reported in a baseline questionnaire and in detailed 24 dietary recall questionnaires in a subset.

**Results:** Genetic risk scores including two and eight single nucleotide polymorphisms (SNPs) explained up to 0.39%, 0.19% and 0.77% of the variance in coffee, tea and combined coffee and tea consumption respectively. A one standard deviation increase in the 8 SNP genetic risk score was associated with a 0.13 cup per day (95% CI: 0.12, 0.14), 0.12 cup per day (95%CI: 0.11, 0.14) and 0.25 cup per day (95% CI: 0.24, 0.27) increase in coffee, tea and combined tea and coffee consumption, respectively. Genetic risk scores also demonstrated positive associations with both caffeinated and decaffeinated coffee and tea consumption. In 48,692 individuals with dietary recall data, the genetic risk scores were positively associated with coffee and tea, (apart from herbal teas) consumption, but did not show clear evidence for positive associations with other beverages. However, there was evidence that the genetic risk scores were associated with lower daily water consumption and lower overall drink consumption.

**Conclusions:** Genetic risk scores created from variants identified in coffee consumption GWAS associate more broadly with caffeinated beverage consumption and also with decaffeinated coffee and tea consumption.

## Introduction

Caffeine, which is most commonly consumed in the form of coffee and tea, is the most widely use psychoactive substance in the world (1). Due to their widespread use, there is considerable interest in the potential effects of coffee, tea and caffeine consumption on health. Whilst much is known about the acute effects of caffeine use (2), less is understood about the health effects of long term use. For example, in observational studies, coffee or caffeine consumption have been linked to lower risk of a number of diseases including cardiovascular disease, diabetes and depression (3-5). However, it is difficult to determine if these relationships are causal, due to the possibility of confounding and reverse causality.

Identification of genetic variants which determine caffeine consumption can help us to better understand the causal effects of use of this substance, by using these variants as instruments for exposures in Mendelian randomisation analyses(6). Unlike measured coffee and tea consumption, genetic variants should be independent of confounders and should not be affected by reverse causality. Recent genomewide association studies (GWAS) have identified eight independent loci that are associated with coffee consumption at the genomewide significance level (7-9). Combining variants together in an allele score has been shown to increase the amount of variance explained in coffee or caffeinated beverage consumption (10, 11). The two most strongly associated loci identified to date are in or near genes which are involved in caffeine metabolism (the cytochrome P450 1A1 and 1A2 (CYP1A1/2) gene region and aryl-hydrocarbon receptor gene (AHR)) (12). *CYP1A2* is the enzyme primarily responsible for metabolising caffeine and *AHR* affects *CYP1A2* activity (12). It is likely that theses variants affect coffee consumption through altering rate of caffeine metabolism; there is evidence that the coffee increasing alleles of these SNPs decrease blood caffeine levels (13, 14). Of note, five of the six additional loci (or close proxies) (in or near the following genes: *GCKR, ABCG2, MLXIPL, BDNF, EFCAB5)* have also been identified in GWAS of other phenotypes, such as body mass index and smoking initiation (9, 15, 16), suggesting that these variants are not specific to coffee or caffeine consumption.

To be suitable instruments for Mendelian randomisation analyses, genetic risk scores for coffee or caffeine consumption should 1) be robustly associated with the exposure of interest, 2) not be associated with potential confounding factors of the exposure-outcome relationship, and 3) only be associated with the outcome through the exposure of interest (i.e., no pleiotropic effects) (17) (although methods for accounting for potential bias due to pleiotropy, such as MR Egger (18), have been developed). As GWAS studies have to date focused on coffee consumption as the phenotypic measure of interest, we sought to explore the associations of these risk scores with consumption of coffee, tea and a broader range of beverages in 114,316 individuals from the UK Biobank study. Although individual associations of SNPs in the 8 genomewide significant loci with coffee and tea have been reported previously for UK Biobank (14), data on their use in genetic risk scores has not been presented. More accurate characterisation of the phenotypes that these genetic risk scores capture is important for interpreting the results of studies that use these variants to assess how coffee, tea or caffeine may affect health outcomes.

## Methods

### Study population

The UK Biobank (www.ukbiobank.ac.uk) recruited over 500,000 men and women (aged 37 to 73 years) between 2006 and 2010 (19). Participants attended one of the 21 assessment centres in England, Wales and Scotland, where they provided information on demographic, lifestyle factors and medical history through interviews and questionnaires and had physical measurements and blood, urine and saliva samples taken. The full protocol for the study is available online: www.ukbiobank.ac.uk/wp-content/uploads/2011/11/UK-Biobank-Protocol.pdf. The UK Biobank study was approved by the North West Multi-Centre Research Ethics Committee and all participants provided written informed consent to participate in the UK Biobank study.

### Genetic risk scores

We created genetic risk scores for caffeine consumption using 8 SNPs from 8 independent loci that reached genomewide significance with coffee consumption in the Coffee and Caffeine Genetics Consortium (CCGC) (rs4410790, rs2470893, rs1260326, rs1481012, rs7800944, rs9902453, rs17685, rs6265) (9). We also created a genetic risk score using just variants in the strongest two loci, *AHR* (rs4410790) and *CYP1A1/2* (rs2470893), which have been identified in multiple GWAS studies of coffee consumption (7, 8). Full details of these SNPs are provided in supplementary methods. Details of the imputation quality for each SNP are shown in Table S1. Unweighted risk scores were created by adding together the number of coffee consumption increasing alleles. Weighted risk scores were created by multiplying each coffee consumption increasing allele by the magnitude of its association with coffee consumption in individuals of European ancestry in the discovery sample of the CCGC genomewide association study (9) (see Table S1 for effect sizes).

### Tea, coffee and other beverage consumption

Information about usual intake of tea and coffee were assessed as part of the baseline questionnaire, which was administered to all participants during their visit to the initial assessment centre. All UK Biobank participants were asked how many glasses of water they drank each day. For each type of drink, answers were provided on a continuous scale and we excluded individuals reporting drinking >25 cups/glasses a day.

Information on tea, coffee and other beverage consumption was also obtained for a subset of participants from a 24 hour diet recall, which had questions on consumption of about 200 commonly consumed items. This questionnaire has been validated previously (20). The questionnaire was administered at the last station of the assessment centre visit towards the end of recruitment and also emailed to all participants with known email addresses (N = 320,000) on four separate occasions between February 2011 and April 2012. For full details see: http://biobank.ctsu.ox.ac.uk/crystal/docs/DietWebQ.pdf. Participants were asked how many times they consumed specific drinks on the previous day. For each type of drink, they could respond with the following options: None, ½, 1, 2, 3, 4, 5, 6+. Full details of the questions asked, the coding of drink consumption and the coding of caffeinated and decaffeinated coffee and tea consumption are provided in supplementary material.

### Statistical analysis

Analyses were conducted in Stata (version 14.2). Associations between genetic risk scores and consumption of each type of beverage were assessed using linear regression, adjusting for age (as a continuous variable), sex and 15 genetic principal components. Robust standard errors were calculated to account for non-normality of residuals. Weighted risk scores were converted to z-scores, so associations are per standard deviation increase. Associations for unweighted risk scores are per additional coffee consumption increasing allele. Primary analyses in the full UK Biobank sample were conducted in all individuals (consumers and non-consumers), but we conducted sensitivity analyses restricted to consumers. Primary analyses in the subset with dietary data were restricted to consumers of each beverage, as many of the beverages were only consumed by a small proportion of the sample. We also tested for associations with any vs no tea and coffee consumption in the full sample using logistic regression. As smoking is known to increase caffeine metabolism through induction of CYP1A2 (21), we performed an additional sensitivity analysis in the full sample stratified by smoking status (never, former and current).

## Results

### Description of study population

A total of 114,316 unrelated individuals of European ancestry were included in the analysis of coffee and tea captured at the initial UK Biobank assessment centre. Dietary recall data was available for a subset of 48,692 of these individuals (Table 1). Males made up just under half of these samples and the mean age was 56.9 and 56.5 years in the full sample and subset respectively. Prevalence of tea and coffee consumption were high (85% of individuals were tea consumers and 78% coffee consumers in the full sample). Median coffee consumption was 2 cups per day (IQR 0.5, 3). Median tea consumption was 3 cups per day (IQR 1,5). Individuals in the dietary recall subset were more likely to have a degree or professional qualification and less likely to be smokers than those in the full sample.

**Table 1.** Description of study samples.

|  | Full sample<br>(N= 114,316) |  | Subset with dietary recall<br>data (N=48,692) |  |
| --- | --- | --- | --- | --- |
| Male: N (%) | 54,024 | 47.26 | 22,789 | 46.80 |
| Age: Mean (SD) | 56.9 | 7.92 | 56.49 | 7.79 |
| **Education** |  |  |  |  |
| None | 20,615 | 18.20 | 4,559 | 9.40 |
| NVQ/CSE/A-levels | 40,898 | 36.10 | 16,578 | 34.17 |
| Degree/professional | 51,770 | 45.70 | 27,377 | 56.43 |
| Any tea consumption | 96,765 | 84.7 | 41,324 | 84.98 |
| Tea (cups per day) | 3 | 1,5 | 3 | 1,5 |
| Any coffee consumption | 90,005 | 78.73 | 39,298 | 80.77 |
| Coffee (cups per day) | 2 | 0.5,3 | 2 | 0.5,3 |
| **Smoking** |  |  |  |  |
| Never | 61,179 | 53.65 | 27,389 | 56.35 |
| Former | 39,006 | 34.21 | 16,848 | 34.66 |
| Current | 13,844 | 12.14 | 4,371 | 8.99 |
| Questionnaires completed:<br>N(%) |  |  |  |  |
| 1 |  |  | 19,093 | 39.21 |
| 2 |  |  | 11,160 | 22.92 |
| 3 |  |  | 9,895 | 20.32 |
| 4 |  |  | 7,198 | 14.78 |
| 5 |  |  | 1,346 | 2.76 |

In the first release of the UK Biobank sample with genetic data, both the 2 SNP and 8 SNP genetic risk scores were associated with coffee and tea intake (see Table 2). The risk scores explained up to 0.39% of the variance in coffee consumed per day, up to 0.19% of the variance in tea consumed per day, and up to 0.77% of the variance in tea and coffee combined. The weighted 8 SNP explained the highest proportion of the variance for both tea and coffee. The unweighted and weight 2 SNP scores performed similarly in terms of variance explained. There was no strong evidence that these associations differed in never, former, or current smokers, although point estimates for the 2 SNP score were generally lower in current smokers and there was some evidence of heterogeneity between smoking groups in the combined tea and coffee analysis (I^2^ = 63%) (Figure S2).

**Table 2.** Associations of 8 and 2 SNP genetic risk scores with tea and coffee consumption (cups per day).

|  | N | Unweighted<sup>1</sup> | R-squared<sup>3</sup> | Weighted<sup>2</sup> | R-squared<sup>3</sup> |
| --- | --- | --- | --- | --- | --- |
| **Cups of coffee per day (including decaffeinated)** |  |  |  |  |  |
| 2 SNP score | 114,316 | 0.12 (0.11, 0.13) | 0.29% | 0.11 (0.10, 0.13) | 0.29% |
| 8 SNP score | 111,645 | 0.063 (0.056, 0.070) | 0.29% | 0.13 (0.12, 0.14) | 0.39% |
| **Cups of tea per day** |  |  |  |  |  |
| 2 SNP score | 114,316 | 0.12 (0.10, 0.13) | 0.16% | 0.11 (0.10, 0.13) | 0.16% |
| 8 SNP score | 111,469 | 0.057 (0.047, 0.066) | 0.13% | 0.12 (0.11, 0.14) | 0.19% |
| **Cups of tea and coffee per day** |  |  |  |  |  |
| 2 SNP score | 114,316 | 0.24 (0.22, 0.26) | 0.61% | 0.23 (0.21,0.24) | 0.61% |
| 8 SNP score | 111,469 | 0.120 (0.110, 0.129) | 0.54% | 0.25 (0.24,0.27) | 0.77% |

Analyses include consumers and non-consumers. 8 SNP score had missing genotype information for 2,847 individuals.
1. Associations per coffee consumption increasing allele, adjusted for age, sex and genetic principal components 2. Associations are per SD increase in genetic risk score, adjusted for age, sex and genetic principal components 3. Calculated from residuals of risk score on genetic principal components, regressed on coffee/tea/tea and coffee per day.

There was evidence that the caffeine genetic risk scores were associated with higher odds of drinking any coffee compared to no coffee (OR per SD increase in 8 SNP weighted genetic risk score: 1.06, 95% CI: 1.05, 1.08). However, there was no clear evidence that the genetic risk scores were associated with consuming either any tea or any tea or coffee combined (Table S3).

### Associations of genetic risk scores by type of drink

In analyses restricted to consumers of each type of drink, in the full sample each additional coffee consumption increasing allele of the 2 SNP genetic risk score was associated with a 0.12 increase in the number of cups of coffee consumed per day (95% CI: 0.10, 0.13), a 0.14 increase in the number of cups of tea consumed per day (95% CI: 0.13, 0.16) and a 0.24 increase in the number of cups of tea and coffee combined (95% CI: 0.22, 0.26). We observed a negative association with water consumption; each additional coffee consumption increasing allele was associated with consuming 0.06 fewer glasses of water per day (95% CI: −0.07, −0.04).

Similar patterns were observed in the subset of individuals with dietary recall data, the 2 SNP unweighted genetic risk score was positively associated with most types of coffee consumption (Figure 1). Each additional caffeine consumption increasing allele was associated with higher consumption of instant, filter, latte, decaffeinated and total coffee. For cappuccino and other coffee, point estimates were in the positive direction but confidence intervals overlapped the null. The genetic risk score was also associated with increased standard tea but not with rooibos, green tea or herbal tea. As observed in the full sample, the genetic risk score was negatively associated with water consumption; each additional coffee consumption increasing allele was associated with a 0.04 decrease (95% CI: −0.06, − 0.02) in the number of glasses of water consumed per day. There was no clear evidence that the genetic risk score was associated with consumption of any of the other non-tea or coffee beverages. The genetic risk score was positively associated with combined coffee and tea consumption (0.18 additional portions consumed per additional coffee consumption increasing allele, 95% CI: 0.16, 0.19), but negatively associated with non-coffee and tea consumption (0.08 fewer portions consumed per additional coffee consumption increasing allele, 95% CI: −0.10, −0.05). Overall, each additional caffeine consumption increasing allele was associated with consumption of 0.10 additional drinks per day (95% CI: 0.07, 0.12).

**Figure 1.**
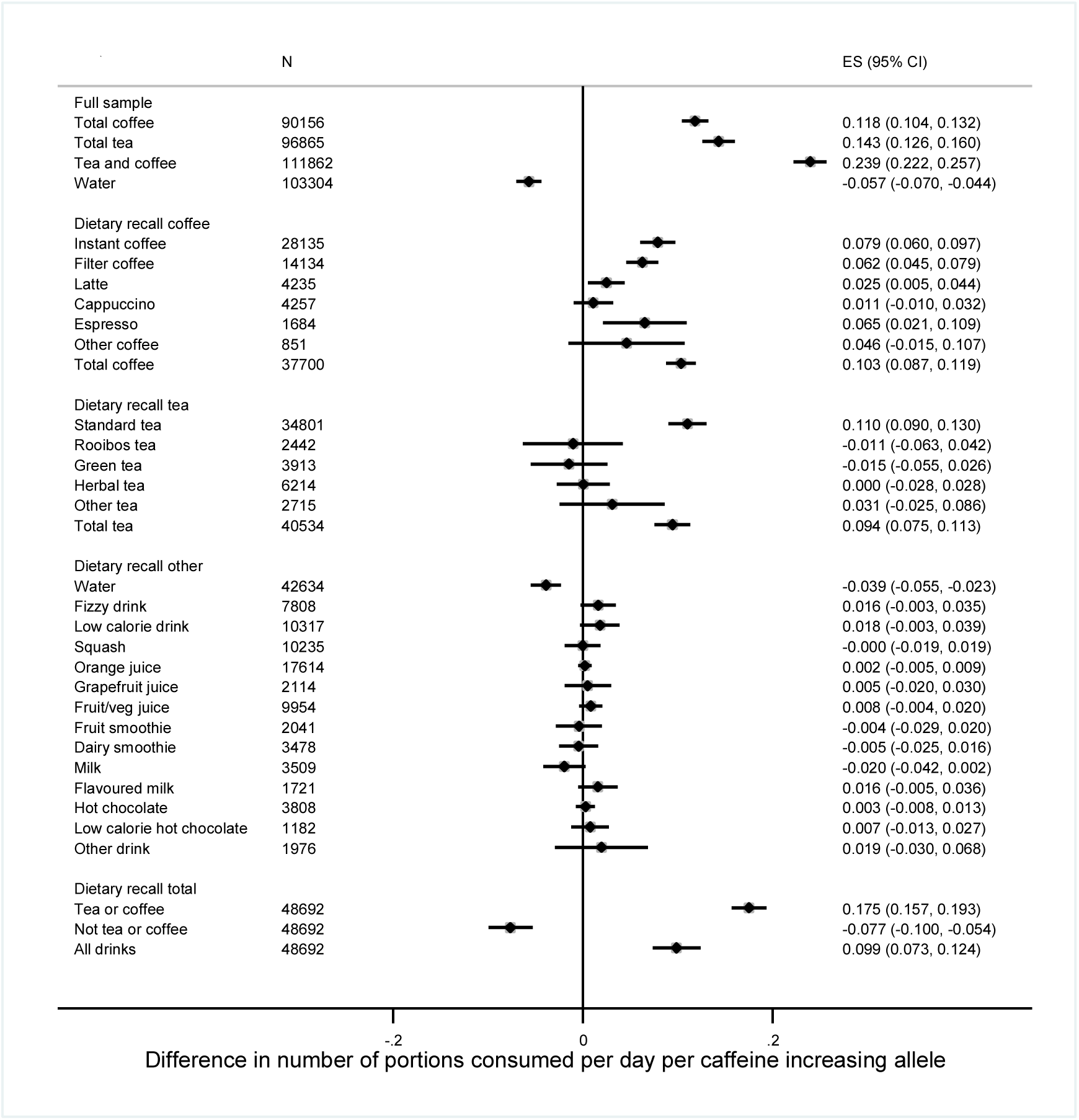
Associations between the 2 SNP genetic risk score and types of drink in the full sample and dietary recall subset. Analyses restricted to consumers of each drink apart from the dietary recall total which included all respondents. Adjusted for age, sex and principal genetic components.

When analyses were repeated using the 8 SNP weighted genetic risk score results were similar (Figure S1). As the initial GWAS release of the UK Biobank contains data from a nested case control study participants were selected on the basis of lung function and smoking status (UK Bileve) (22), we also repeated analyses excluding these individuals. This made little difference to the observed associations (data not shown).

### Caffeinated and decaffeinated tea and coffee consumption

The 2 SNP genetic risk score was positively associated with both caffeinated and decaffeinated tea and coffee consumption (Figure 2). Point estimates for decaffeinated coffee consumption and caffeinated coffee consumption were similar in the full UK Biobank sample, but were smaller for decaffeinated coffee consumption (beta 0.06, 95% CI: 0.03, 0.09) than for caffeinated consumption (beta 0.09, 95% CI: 0.08, 0.11) in the dietary recall sample. For tea consumption, point estimates were similar for both caffeinated and decaffeinated in the dietary recall study. Similar patterns were observed for the 8 SNP score (see Figure S4).

**Figure 2.**
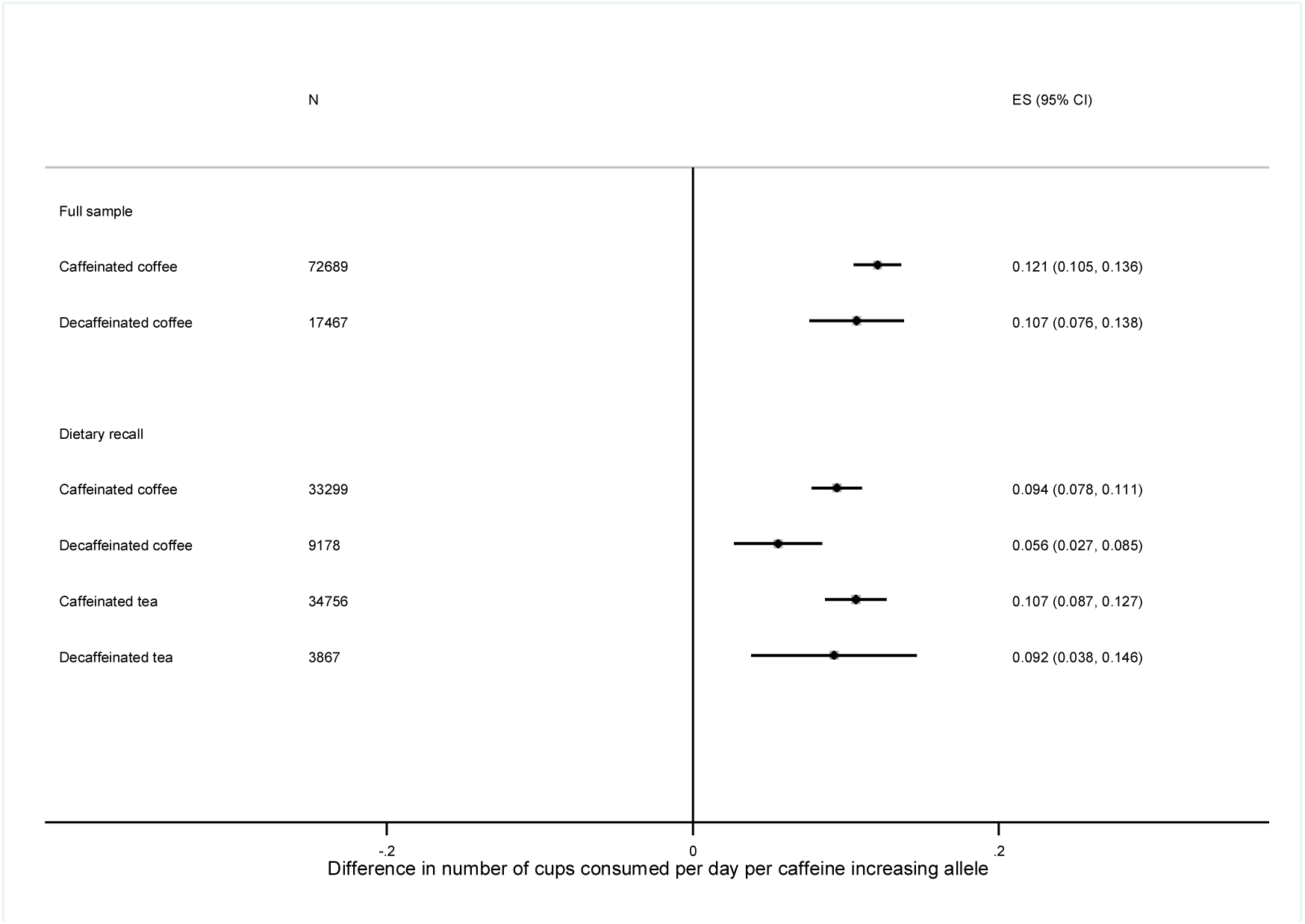
Associations between the 2 SNP genetic risk score and caffeinated and decaffeinated coffee. Analyses restricted to consumers of each drink. Adjusted for age, sex and principal genetic components.

### Association of genetic risk scores with demographic factors

There was no clear evidence that the genetic risk scores were associated with age, education levels, income or level of deprivation (Supplementary Table S3). However, there was some evidence for a positive association with alcohol consumption (OR for daily vs non daily consumption per SD increase in 8 SNP genetic risk score 1.02 (95%CI, 1.01, 1.04), p=0.002. The 2 SNP genetic risk scores also showed suggestive evidence of being associated with current smoking (OR for current smoking per coffee increasing allele 0.98 (95% CI: 0.96, 1.00) and sex (OR for female sex per coffee increasing allele 0.99 (95% CI: 0.97, 1.00), although evidence for these associations was weaker with the 8 SNP genetic risk score.

## Discussion

We have confirmed associations of genetic risk scores for coffee consumption using variants identified in published GWAS (7-9), with consumption of both coffee and tea in a large sample of older adults of European ancestry from the UK. Using a more detailed measure of beverage consumption, the 24 hour dietary recall, we found that the genetic risk scores were associated with most types of coffee consumption, and with black tea but not herbal tea consumption. There was no clear evidence that the genetic risk scores increased consumption of other types of nonalcoholic beverages, but they were associated with decreased consumption of water and total drinks excluding tea and coffee, suggesting that coffee and tea drinks are consumed instead of rather than in addition to other types of drinks, consistent with fluid homeostasis.

Associations of the genetic risk scores with both coffee and tea support the use of coffee genetic risk scores as determinants of caffeinated beverage consumption (and likely caffeine consumption in general) rather than as specific to coffee consumption. Variance explained in combined coffee and tea consumption was 0.54-0.77% compared to 0.13-0.39% for coffee or tea alone. This has been shown previously in a study of women from the UK (11) and is unsurprising given that the two strongest coffee-related variants in the risk score (in *AHR* and *CYP1A1/2)* are in or near genes in caffeine metabolising pathways. Although we were unable to assess associations with other specific caffeinated beverages (e.g., cola), we found suggestive evidence for positive associations with carbonated and low calorie drinks. Coffee tends to have a higher caffeine content than tea and the genetic risk scores explain more of the variance in coffee than in tea consumption. In addition, the genetic risk scores determined whether individuals were coffee consumers but not whether they were tea consumers. Given the variation in consumption of different types of caffeinated drink consumption between countries (23) and by demographic factors (e.g., age) (12), it is likely that these associations will be context specific. Whilst these genetic risk scores may serve as instruments for tea as well as coffee consumption in Mendelian randomisation analyses in UK populations, they may not associate with tea consumption in countries with low levels of tea consumption.

The effect sizes we observed with coffee consumption were similar to those observed in previous GWAS of coffee consumption (8, 9). Each coffee consumption increasing allele of the SNPs in *AHR* and *CYP1A1/2* was associated with an increase of about 0.1 cups of coffee. Our data suggest that using the 8 SNP weighted genetic risk score increases power to detect associations with caffeinated drink consumption over the 2 SNP genetic risk score (weighted or unweighted), but only by a relatively small amount (up to a 34% increase). Given that the other genetic variants are known to be associated with other non-caffeine related phenotypes, use of the 2 SNP risk score as an instrument for coffee or caffeine consumption may be more appropriate for some Mendelian randomisation analyses and is unlikely to result in substantial loss of power. However, it is also important to note that CYP1A2 metabolises xenobiotic substrates other than caffeine (24), so SNPs in *CYP1A1/2* and *AHR* may have downstream effects which do not act through caffeine.

The genetic risk scores associated with both caffeinated and decaffeinated coffee and tea consumption in UK Biobank. Associations of *AHR* variants with decaffeinated coffee consumption in the same direction as for caffeinated coffee consumption were also reported in the CCGC GWAS (9). This is likely to be explained by a continuation of coffee and tea drinking behaviour in individuals who have switched from caffeinated to decaffeinated coffee and tea consumption. We did not observe associations with herbal, green or rooibos tea in UK Biobank; it is possible that these drinks are not consumed in high enough quantities or that they are not direct replacements for caffeinated beverages. Importantly, it is likely that there is some misclassification of decaffeinated tea and coffee consumption due to the way individuals were asked about decaffeinated drink consumption in UK Biobank (only being able to say they drank all decaffeinated, all caffeinated or a mixture). Therefore, some caution must be exercised when interpreting these results.

To our knowledge, the negative association between these genetic variants and water consumption has not been reported previously. The negative association was also observed for overall non-tea and non-coffee drink consumption. We think that the most parsimonious explanation for this finding is that this is a downstream effect of increased coffee and tea consumption; individuals drinking more coffee and tea consume fewer other drinks. We cannot completely rule out the diuretic effect of caffeine as an explanation for this finding. There is evidence that the coffee consumption increasing alleles in *AHR, CYP1A1/2* are associated with reduced blood caffeine levels as caffeine metabolism is increased in these individuals (13). Given that caffeine is a known diuretic (25), lower caffeine levels could in theory result in higher hydration and lower consumption of non-caffeinated beverages. However, there is evidence that normal levels of caffeine consumption via tea and coffee are too low to result in net fluid loss and dehydration (25).

Whilst these genetic risk scores appear to be robust instruments for caffeinated beverage consumption, we found some evidence within UK Biobank for associations with other traits, most notably daily alcohol consumption. This could be of concern for use of these risk scores as proxies for coffee or caffeine consumption in Mendelian randomisation studies as this would potentially violate the assumption of no pleiotropy. A previous Mendelian randomisation study using variants in *CYP1A1/2* and *AHR* did not find clear evidence for associations with potential confounders (26), but did not investigate alcohol consumption. Further work is required to determine whether an association between the coffee/caffeine related genetic risk scores represents a true association and if so, whether this effect is direct (not through caffeine consumption) or indirect (representing a causal effect of caffeine on alcohol consumption).

There are several limitations to this analysis which should be considered. Firstly, these analyses have been conducted in individuals of European ancestry and may not be generalizable to other ethnicities. As discussed above, variation in type of caffeine consumption by population may also limit generalisability of these results. Secondly, the UK Biobank had a low response rate (around 5%) and is not likely to be representative of individuals of this age group in the UK (27). Additionally, selection into the UK BiLEVE GWAS sample make this sample unrepresentative of UK Biobank as a whole. However, exclusion of individuals in the BiLEVE sample made little difference to results. Thirdly, there is likely to be measurement error in the reporting of beverage consumption. In the main questionnaire, individuals could report consumption of tea and coffee as exact number of cups consumed per day but in the dietary recall study, individuals could only report consumption up to a maximum of 6+ portions per day. Although patterns of association were very similar for tea and coffee between the two, this restriction of the maximum value is likely to impact upon magnitude of associations, particularly when summing across categories. Therefore, effect sizes should be interpreted with caution. Finally, this study does not assess the association of these risk scores with blood caffeine levels. As discussed above, variants in *CYP1A1/2* and *AHR* which increase coffee consumption decrease plasma caffeine (13, 14). A recent analysis including data from UK Biobank has also demonstrated that the SNPs in *GCKR, EFCAB5, POR* also associate in opposing directions with coffee consumption and plasma caffeine metabolites (14).

In conclusion, genetic risk scores compiled using variants from coffee consumption GWAS appear to associate more broadly with caffeine containing beverages (or decaffeinated versions of these beverages). This is an important consideration for using these scores as instruments for coffee, tea or caffeinated beverage consumption in Mendelian randomisation analyses. What these variants relate to within each specific analysis sample will be important for interpreting results of Mendelian randomisation analyses. Finally, association of these genetic risk scores with non-caffeine related phenotypes e.g., alcohol consumption may invalidate their use to assess downstream health effects of caffeine consumption.

## Funding and Acknowledgements

UK Biobank: This research has been conducted using the UK Biobank Resource. AET and MRM are members of the UK Centre for Tobacco and Alcohol Studies, a UKCRC Public Health Research: Centre of Excellence. Funding from British Heart Foundation, Cancer Research UK, Economic and Social Research Council, Medical Research Council, and the National Institute for Health Research, under the auspices of the UK Clinical Research Collaboration, is gratefully acknowledged. This work was supported by the Medical Research Council (grant number: MC_UU_12013/6).

## Conflicts of interest

The authors have no conflicts of interest to declare.

## Declarations of interest

None

